# The phototrophic bacteria *Rhodomicrobium* spp. are novel chassis for bioplastic production

**DOI:** 10.1101/2023.05.17.541187

**Authors:** Eric M. Conners, Karthikeyan Rengasamy, Arpita Bose

## Abstract

Polyhydroxybutyrate (PHB) is a bio-based, biodegradable alternative to petroleum-based plastics. PHB production at industrial scales remains infeasible, in part due to insufficient yields and high costs. Addressing these challenges requires identifying novel biological chassis for PHB production and modifying known biological chassis to enhance production using sustainable, renewable inputs. Here, we take the former approach and present the first description of PHB production by two prosthecate photosynthetic purple non-sulfur bacteria (PNSB), *Rhodomicrobium vannielii* and *Rhodomicrobium udaipurense.* We show that both species produce PHB across photoheterotrophic, photoautotrophic, photoferrotrophic, and photoelectrotrophic growth conditions. Both species show the greatest PHB titers during photoheterotrophic growth on butyrate with dinitrogen gas as a nitrogen source (up to 44.08 mg/L), while photoelectrotrophic growth demonstrated the lowest titers (up to 0.13 mg/L). These titers are both greater (photoheterotrophy) and less (photoelectrotrophy) than those observed previously in a related PNSB, *Rhodopseudomonas palustris* TIE-1. On the other hand, we observe the highest electron yields during photoautotrophic growth with hydrogen gas or ferrous iron electron donors, and these electron yields were generally greater than those observed previously in TIE-1. These data suggest that non model organisms like *Rhodomicrobium* should be explored for sustainable PHB production and highlights utility in exploring novel biological chassis.

## Introduction

Plastics are an inescapable part of modern life. They are ubiquitous throughout industries like transportation, food, health-care, and energy, with packaging as the single largest market (Geyer et al., 2017; Rosenboom et al., 2022). Many plastics are single-use, and no commonly used plastics are biodegradable (Geyer et al., 2017). Consequently, global plastic waste is immense: from 1950 - 2015, we produced 6300 million metric tons of plastic waste, 79% of which accumulated in landfills or the environment (Geyer et al., 2017). Exacerbating this pollutive pipeline, conventional plastics are made from fossil fuels, with each stage of the plastic lifecycle contributing to greenhouse gas emissions. By 2050, plastic production will be responsible for almost 13% of the global carbon budget, equal to the combined emissions of 615 coal-fired power stations (*Plastic & Climate - The Hidden Costs of a Plastic Planet*, 2019). One way to address these challenges is to reimagine plastics as part of a circular economy. In this scenario, plastic production would rely on renewable energy and materials, and plastic waste would be eliminated via sustainable degradation practices and/or the reuse and recycling of plastic to support other economic processes. To accomplish these goals, we need novel polymers that fit these criteria.

The term “bioplastic” is invoked when discussing a range of materials with distinct origins and end of life options. Bioplastics may originate from biomass or be biodegradable. But not all bio-based materials are biodegradable (e.g., bio-based polyethylene) and some biodegradable plastics are made using fossil fuels (e.g., polybutylene succinate) (Rosenboom et al., 2022). Bioplastics that are both bio-based and biodegradable can address the shortcomings associated with petrochemical plastic lifecycle and integrate within a circular economy (Rosenboom et al., 2022). One such class of polymers is polyhydroxyalkanoates (PHAs). The most studied PHA, polyhydroxybutyrate (PHB), has several properties that make it suitable as a bio-based plastic: it is thermoplastic, water insoluble, moldable, and biodegradable (Dobrogojski et al., 2018; Sehgal and Gupta, 2020). Furthermore, PHB is quantitatively comparable or superior to petrochemical based and bio-based polymers across several dimensions, including tensile strength (20-40 MPa), crystallinity (50-60%), and melting temperature (165-175°C) (McAdam et al., 2020).

PHB is produced by diverse microbes as intracellular carbon and energy stores. PHB producers comprise species from as many as 75 different genera including *Ralstonia eutropha* (or *Cupriavidus necator*), *Rhodopseudomonas palustris* TIE-1, as well as undefined cultures enriched from environmental or industrial sources (e.g., activated sludge from wastewater) (Ranaivoarisoa et al., 2019; Riedel et al., 2015; Sen et al., 2019; Sindhu et al., 2011). While the available chassis for PHB production has expanded, their commercial utility remains questionable due to insufficient yield, reliance on energy- or cost-intensive inputs, and narrow metabolisms (McAdam et al., 2020). These shortcomings can be addressed via modifications to growth conditions and genetic manipulations; however, novel chassis may offer improved production while providing an opportunity to explore the mechanisms and physiological role of intracellular PHB accumulation.

Environmental bacteria with diverse metabolisms provide a unique window into PHB production. A recent study showed that *Rhodopseudomonas palustris* TIE-1 – also a phototrophic purple non-sulfur bacterium – produces PHB across various phototrophic growth conditions (Ranaivoarisoa et al., 2019). The authors found PHB yields as low as 1.57 mg/L when TIE-1 was grown on succinate with ammonium chloride (NH_4_Cl) as a nitrogen source, and up to 16.57 mg/L when grown on butyrate in nitrogen (N_2_)-fixing conditions. While this suggests that photoheterotrophy using a highly reduced substrate like butyrate is the preferred condition with respect to total yield, the authors showed that photoferrotrophy (i.e., photosynthetic iron oxidation) resulted in the highest specific productivity (PHB mg/L/cell/h). Additionally, the authors assessed PHB production via a microbial electrosynthesis (MES) platform, wherein cells acquire electrons from a poised electrode via extracellular electron uptake (EEU). While PHB yields during MES were relatively low, TIE-1 demonstrated high electron yield. In summary, these three vastly distinct growth conditions offer strengths and weaknesses associated with product formation. Moreover, the authors observed variation in growth kinetics and carbon yields, suggesting even more complexity.

Distantly related photosynthetic purple non-sulfur bacteria may offer additional insights into the relationship between metabolism and PHB production. The genus *Rhodomicrobium* comprises only a handful of known species, with two primary representatives: *R. vannielii* and *R. udaipurense.* They are microaerobic to anaerobic bacteria found in freshwater soils and sediments, particularly among plant microbiomes (Town et al., 2023; Wei et al., 2020; Zuo et al., 2022). They are characterized by their polymorphic lifecycle comprising stalk-forming cells and motile daughter cells (Duchow and Douglas, 1949; Whittenbury and Dow, 1977). Both species are capable of nitrogen fixation as well as chemoheterotrophic, photoheterotrophic, and photoautotrophic metabolisms, including photoferrotrophy (Duchow and Douglas, 1949; Ehrenreich and Widdel, 1994; Whittenbury and Dow, 1977). More recently, *R. vannielii* and *R. udaipurense* were shown to conduct extracellular electron uptake (EEU), wherein outer membrane protein complexes facilitate electron uptake from solid phase conductive substances (e.g., poised electrodes) (Gupta et al., 2019). This makes them appealing for bioproduction studies, as it affords the opportunity to explore the relationship between metabolism and product formation while identifying pros and cons associated with each condition.

In the present study, we cultivate *R. vannielii* and *R. udaipurense* in anoxic conditions under varying carbon, electron, and nitrogen regimes. We quantify PHB production across several parameters and calculate the PHB carbon and electron yields. We also examine possible relationships between carbon yield, electron yield, and PHB production.

## Materials and Methods

### Bacterial strains, media, and growth conditions

*Rhodomicrobium vannielii* ATCC 17100 was purchased from DSMZ (Leibniz Institute, Braunschweig, Germany). *Rhodomicrobium udaipurense* JA643 was acquired from the University of Hyderabad (Hyderabad, India). Both strains were incubated anaerobically in bicarbonate-buffered freshwater (FW) medium (Ehrenreich and Widdel, 1994) prepared under the flow of N_2_-CO_2_ (80%/20%) gas. Either 5.61 mM NH_4_Cl or N_2_ gas (~103 kPa) was provided as a nitrogen source. Cultures were incubated without shaking at a 30-cm distance from a 60-W incandescent light bulb at 30°C, unless otherwise stated. Time-course cell growth was monitored using Spectronic 200 (Thermo Fisher Scientific, USA). All sample manipulations were performed inside an anaerobic chamber with 5% H_2_/75% N_2_/20% CO_2_.

For photoheterotrophic growth, 15 mL of FW medium – supplemented with anoxic sodium butyrate to a final concentration of 10 mM – was dispensed in Balch tubes sealed with sterile butyl stoppers and capped with an aluminum crimp. Cultures pre-grown to an optical density (OD_660_) of 1.0 were washed with FW medium (without NH_4_Cl) and inoculated to a final OD_660_ of 0.01. The resulting cultures were pressurized with N_2_-CO_2_ (80%/20%) to a final headspace pressure of 103 kPa.

For photoautotrophic growth with H_2_ gas as an electron donor, 15 mL of FW medium was prepared in Balch tubes and inoculated as described above. The headspace was filled with H_2_-CO_2_ (80%/20%) to a final pressure of 103 kPa For the N_2_-fixing condition using H_2_ as an electron donor, the headspace was filled to 48 kPa with N_2_-CO_2_ and then to 103 kPa with H_2_-CO_2_. Nitrogen fixing cultures were inoculated to a final OD_660_ of 0.1 to promote cell growth.

For photoautotrophic growth with Fe(II) as an electron donor, approximately 500 mL of FW medium was prepared in a sterile, capped, and crimped serum bottle under the flow of N_2_-CO_2_ (80%/20%) at 103 kPa. The media was then supplemented with anoxic sterile stocks of nitrilotriacetic acid (NTA) to a final concentration of 10 mM and iron chloride (FeCl_2_) to a final concentration of 5 mM. Then, 80 mL of the FW/FeCl_2_/NTA medium was aliquoted to sterile, capped, and crimped serum bottles before inoculation with a pre-grown culture to a starting OD_660_ of 0.1.

### Cell enumeration

*R. vannielii* and *R. udaipurense* cultures grown photoheterotrophically in FW medium supplemented with NH_4_Cl and 10 mM sodium butyrate were harvested at mid-logarithmic phase (OD_660_ ~ 0.5) and diluted or reconcentrated in FW media to generate a range of optical densities within logarithmic phase. The number of cells in each of these cell suspensions was quantified using a Petroff-Hauser chamber (Hausser Scientific, Horsham, PA, USA) according to the manufacturer’s protocol. Cells were visualized and counted using phase contrast microscopy. The OD_660_ of *Rhodomicrobium* spp. vs. cell numbers/mL was plotted to obtain a standard curve.

### Bioelectrochemical growth

Photoelectroautotrophic growth was performed using single-chamber three-electrode configured seal-type bioelectrochemical cells (BECs) (C001 Seal Electrolytic Cell, Xi’an Yima Opto-electrical Technology Com., Ltd, China). The three electrodes were configured as follows: a 5 cm_2_ Pt foil counter electrode, an Ag/AgCl reference electrode submerged in 3 M KCl, and a 1 cm_2_ carbon felt (Fuel Cell Earth, Woburn, Massachusetts, USA) working electrode. Working electrodes were poised at +100 mV vs. standard hydrogen electrode using a multichannel potentiostat (Gamry Instruments, Warmister, PA) that was operated continuously. 75 mL of FW media was dispensed into sterile BECs, which were sealed and made anaerobic by purging and pressurizing to 48 kPa with N_2_-CO_2_ (80%/20%). BECs were connected to the potentiostat and monitored overnight for evidence of residual oxygen presence or any contaminations prior to inoculation. BECs were inoculated with 5 mL of pre-grown cells to a final OD_660_ of 0.2. The resulting cultures were purged and pressurized to 48 kPa with N_2_-CO_2_ (80%/20%) before reattaching to the potentiostat. BECs were operated continuously at 26°C and at a 30-cm distance from a 60-W incandescent light bulb. Cell growth was monitored visually and final OD_660_ readings were taken using Spectronic 200 (Thermo Fisher Scientific, USA).

### Analytical Procedures

#### PHB measurement

10 mL (photoheterotrophy, photoautotrophy with H_2_) or 40-50 mL (photoferrotrophy, photoelectrotrophy) of cell cultures at an OD_660_ of 0.7 (unless stated otherwise) was pelleted at 8000xg for 10 min and stored at −80°C until PHB extraction and analysis were performed. 1 mL of water (LC-MS grade) and 600 µL of methanol (HPLC grade) were added to arrest metabolic activity. 10 mg/mL of poly[(*R*)-3-hydroxybutyric acid] (Sigma-Aldrich, USA) was used as a PHB standard. Extraction of PHB was followed by its conversion to crotonic acid. The concentration of crotonic acid was measured using an Agilent Technologies 6420 Triple Quad LC/MS as follows: using Hypercarb column, particle 5 µm, 100 × 2.1 mm (Thermo Fisher Scientific, USA) as stationary phase; water with 0.1% (v/v) formic acid as phase A; acetonitrile and 1% (v/v) formic acid as phase B. The injection volume was 5 µL; the flow rate was set at 500 µL min_−1_; the column temperature was set at 15 °C and the gas temperature was 300 °C [31]. PHB was detected as crotonic acid with mass to charge ratio (*m*/*z*) = 87 which was normalized to bacterial cell number. Details on PHB extraction, PHB carbon yield, and PHB electron yield calculations are described in supplemental methods.

#### H_2_, N_2_, and CO_2_ measurement

Time-course H_2_, N_2_, and CO_2_ from relevant growth conditions were analyzed using gas chromatography (Shimadzu BID 2010-plus, equipped with Rt_®_-Silica BOND PLOT Column: 30 m × 0.32 mm; Restek, USA) with helium as a carrier gas. At each time point, 10 – 50 µL of gas was sampled from the headspace of the serum bottles using a Hamilton™ gas-tight syringe and injected into the column. To quantify dissolved CO_2_, 1 mL of filtered (using 0.22 µm PES membrane filter) aqueous samples from each reactor were collected and injected into helium-evacuated 12-mL septum-capped glass vials (Exetainer, Labco, Houston, TX, USA) containing 1 mL of 85% phosphoric acid. Then, 10 µL of evolved CO_2_ in the headspace was injected into the column. The total CO_2_ in the reactors was calculated as described previously [https://www.pnas.org/doi/abs/10.1073/pnas.1006175107].

#### Sodium butyrate measurement

Time-course consumption of sodium butyrate was quantified using Ion Chromatography metrohm 881 Compact Pro using a Metrosep organic acid column (250 mm length). 0.5 mM H_2_SO_4_ with 15% acetone was used as eluent at a flow rate of 0.4 mL min_−1_ with suppression (10 mM LiCl regenerant).

#### Fe(II) measurement

Time-course Fe(II) concentration was measured using the Ferrozine Assay as described previously (Bose and Newman, 2011).

### Bioelectrochemical analyses

To assess the ability of *Rhodomicrobium* sps. to conduct electron uptake during photoelectrotrophy, chronoamperometry was conducted for 144 h. The working electrode was poised at +100 mV vs. standard hydrogen electrode (SHE). Current response was assessed using Gamry Echem Analyst software (Gamry Instruments, Warmister, PA). We assessed current response across two parameters: first, we measured current density vs. time, which shows the real-time current response of bacteria-electrode electron transfer. Second, we integrated the current density over time to determine the total current uptake in Coulombs (ampere*second).

### Identifying PHB cycle genes in *Rhodomicrobium* spp

*Rhodomicrobium vannielii* ATCC17100 (JAEMUJ010000000) and *Rhodomicrobium udaipurense* JA643 (JAEMUK010000000) genomes were downloaded from the National Center for Biotechnology Information database. We searched the downloaded genomes for putative PHB cycle genes with RAST’s (https://rast.nmpdr.org/) BLASTp function using the amino acid sequences of known TIE-1 homologs as queries.

## Results

### Rhodomicrobium vannielii and Rhodomicrobium udaipurense encode putative PHB cycle genes

Known PHB cycle genes are encoded by diverse bacteria including the freshwater soil phototroph *Rhodopseudomonas palustris* TIE-1, the plant symbiont *Sinorhizobium meliloti,* and the H_2_-oxidizing chemolithotroph *Cupriavidus necator* (D’Alessio et al., 2017; Ranaivoarisoa et al., 2019; Slater et al., 1988). Before exploring PHB production in *R. vannielii* and *R. udaipurense*, we searched their recently re-sequenced genomes for putative homologues of the PHB cycle genes encoded by *R. palustris* TIE-1 (Conners et al., 2021). Like *Rhodomicrobium*, TIE-1 is in the order Hyphomicrobiales capable of phototrophic growth using diverse carbon, electron, and nitrogen sources (Ranaivoarisoa et al., 2019). We identified homologues in both *R. vannielii* and *R. udaipurense* (**Figure 1**), suggesting they produce PHB using a similar biochemical pathway to TIE-1. Therefore, we used TIE-1’s hypothesized PHB synthesis pathway as a basis for reconstructing the *Rhodomicrobium* PHB synthesis pathway (**Figure 1A**) (Ranaivoarisoa et al., 2019). The putative pathway is as follows: first, β-ketothiolase (PhaA) condenses two acetyl-CoAs into acetoacetyl-CoA. Acetoacetyl-CoA is then reduced to (R)-3-hydroxybutyryl-CoA via acetoacetyl-CoA reductase (PhaB). The PHB polymerase (PhaC) and depolymerase (PhaZ) are responsible for PHB polymerization and depolymerization, respectively. This is crucial to PHB’s role as an intracellular carbon and energy store; when cells call upon PHB during times of nutrient stress, PhaZ catabolizes PHB granules, making them bioavailable to the cell (Quelas J. I. et al., 2016). Furthermore, PhaC expression is controlled by the repressor PhaR in *B. diazoeficiens*, suggesting that it plays a similar role in *Rhodomicrobium* (Quelas J. I. et al., 2016). Finally, the phasin proteins (PhaP) likely dictate the number and size of PHB granules, as in B*. diazoeficiens* (Quelas J. I. et al., 2016). We identified homologs of all PHB synthesis genes in both *Rhodomicrobium* genomes (**Figure 1**). This includes five acetyl-CoA acetyltransferases (*phaA*), one acetoacetyl-CoA reductase (*phaB*), two PHB polymerases (*phaC*), one PHB depolymerase (*phaZ*), and three phasins (*phaR*) (**Figure 1A**). Both strains show similar genetic architecture, with *phaA* and *phaB* forming a putative operon nearby *phaR,* which is encoded on the opposite strand (**Figure 1B**).

**Figure 1.**
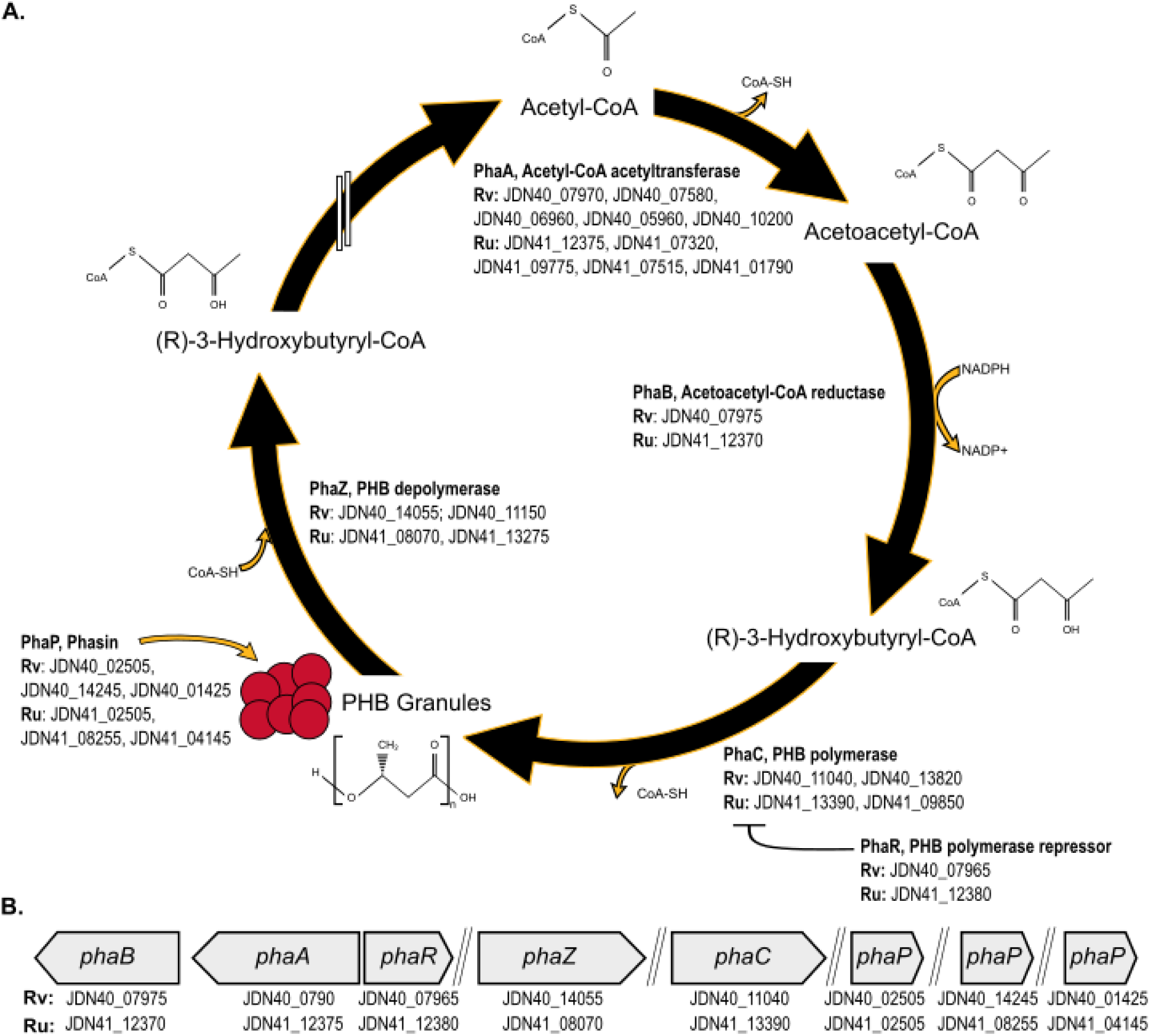
The putative PHB synthesis cycle and genes in R. vannielii and R. udaipurense. **A.** The putative PHB synthesis cycle of R. vannielii (Rv) and R. udaipurense (Ru) based on the hypothesized PHB cycle of Rhodopseudomonas palustris TIE-1 (Ranaivoarisoa et al. 2019). Acetyl-CoA is produced from organic or inorganic carbon sources followed by condensation of two acetyl-CoAs into acetoacetyl-CoA by PhaA. Acetoacetyl-CoA is reduced by the enzyme acetoacetyl-CoA reductase (PhaB) in an NAD dependent manner, producing (R)-3-Hydroxybutyryl-CoA. (R)-3-Hydroxybutyryl-CoA is polymerized into PHB granules by PHB polymerase (PhaC). When cells call upon PHB granules for carbon and/or energy reserves, PHB depolymerase (PhaZ) degrades the granules back into (R)-3-Hydroxybutyryl-CoA and then to Acetyl-CoA via multiple enzymatic reactions (depicted as double white lines). PhaR represses the expression of PhaC, while phasins (PhaP) dictate size and number of PHB granules. Locus tags for each strains’ homologs are listed under their respective genes. **B**. The putative genes involved in the PHB synthesis cycle of Rv and Ru based on highest degree of similarity to TIE-1 homologs. PHB: Polyhydroxybutyrate. CoA-SH: Coenzyme A. NADPH: Nicotinamide adenine dinucleotide phosphate. Adapted from Ranaivoarisoa et al. (2019).

### *Rhodomicrobium* shows variable growth dynamics across different photoautotrophic and photoheterotrophic conditions

We characterized *Rhodomicrobium* growth across several anoxic growth conditions with distinct carbon, electron, and nitrogen sources. These included photoautotrophy (H_2_ electron donor, CO_2_ carbon source), photoheterotrophy (sodium butyrate electron and carbon source), and photoferrotrophy (FeCl_2_ electron donor, CO_2_ carbon source). All growth conditions included light as a source of energy. We generally saw faster generation times during photoheterotrophy, ranging from 15.2 ± 2.3 to 25.5 ± 2.3 hours (**Table 1**). Photoautotrophy demonstrated generation times ranging from 26.9 ± 0.47 to 61.4 ± 6.3 hours (**Table 1**). Photoferrotrophy was among the slowest and most variable growth conditions, with generation times from 120.3 ± 23.2 to 464.4 ± 270.9 (**Table 1**).

**Table 1.** Growth metrics for *Rhodomicrobium* species under different carbon, electron, and nitrogen regimes

| Strain | Carbon Source/Electron Donor/Nitrogen Source | Lag time (h) | Generation time (h) | P-value |
| --- | --- | --- | --- | --- |
| *R. vannielii* | CO<sub>2</sub> / H<sub>2</sub> / N<sub>2</sub> | 20 | 53.5 (2.5) | **2.34E-04** |
|  | CO<sub>2</sub> / H<sub>2</sub> / NH<sub>4</sub>Cl | 12 | 26.9 (0.47) |  |
|  | Butyrate / N<sub>2</sub> | 20.2 | 25.5 (2.3) | **1.07E-02** |
|  | Butyrate / NH<sub>4</sub>Cl | 12.2 | 15.2 (2.3) |  |
|  | CO<sub>2</sub> / FeCl<sub>2</sub> / N<sub>2</sub> | 40 | 342.1 (20.5) | 5.22E-02 |
|  | CO<sub>2</sub> / FeCl<sub>2</sub> / NH<sub>4</sub>Cl | 34.5 | 464.4 (270.9) |  |
|  | CO<sub>2</sub> / CF Cathode / N<sub>2</sub> | 70.1 | 140.3 (24.8) | 2.50E-01 |
|  | CO<sub>2</sub> / CF Cathode / NH<sub>4</sub>Cl | 41.8 | 83.6 (7.7) |  |
| *R. udaipurensis* | CO<sub>2</sub> / H<sub>2</sub> / N<sub>2</sub> | 20 | 61.4 (6.3) | **1.69E-03** |
|  | CO<sub>2</sub> / H<sub>2</sub> / NH<sub>4</sub>Cl | 12 | 27.4 (1.4) |  |
|  | Butyrate / N<sub>2</sub> | 20.2 | 17.5 (0.3) | 2.55E-01 |
|  | Butyrate / NH<sub>4</sub>Cl | 12.2 | 15.9 (1.2) |  |
|  | CO<sub>2</sub> / FeCl<sub>2</sub> / N<sub>2</sub> | 34.5 | 401.0 (42.3) | **3.15E-04** |
|  | CO<sub>2</sub> / FeCl<sub>2</sub> / NH<sub>4</sub>Cl | 22 | 120.3 (23.2) |  |
|  | CO<sub>2</sub> / CF Cathode / N<sub>2</sub> | 52.1 | 117.1 (17.2) | 4.05E-1 |
|  | CO<sub>2</sub> / CF Cathode / NH<sub>4</sub>Cl | 58.6 | 104.3 (5.8) |  |

See *Materials and Methods* for description of lag time and generation time calculations. Lag time is defined as half of the time required to reach the first doubling based on growth curves. For growth on CF cathode, lag time is half of the generation time because cells immediately enter exponential phase and only initial and final optical densities were measured. CF: carbon felt. N<sub>2</sub>: nitrogen-fixing growth with dinitrogen gas as sole nitrogen source. NH<sub>4</sub>Cl: non-nitrogen fixing growth with ammonium chloride as sole nitrogen source. *P*-values are calculated between N<sub>2</sub> and NH<sub>4</sub>Cl generation times for each respective condition. Significant *P*-values are bolded. () = standard deviation values from N ≥ 3 except for *R. vannielii* CO<sub>2</sub>/CF Cathode/N<sub>2</sub> (N = 2).

We also examined the ability of *Rhodomicrobium spp*. to fix nitrogen. Both strains were able to grow in N_2_-fixing conditions, though generation times were usually slower. For example, both strains demonstrated slower growth during N_2_-fixing photoautotrophy, with *R. vannielii’s* generation time slowing to 53.5 hours (*p* = 2.34E-04) and *R. udaipurense’s* generation time slowing to 61.4 hours (*p* = 1.69E-03) (**Table 1**). Likewise, *R. vannielii* showed a significantly slower growth rate during N_2_-fixing photoheterotrophy compared to non-fixing (*p* = 1.07E-02), while *R. udaipurense* showed a significantly slower growth rate during N_2_-fixing photoferrotrophy (*p =* 3.15E-04) (**Table 1**). While we observed slower generation times during other N_2_-fixing growth experiments, these were not statistically significant compared to their non-fixing counterparts. Overall, these data show that *Rhodomicrobium* can grow in diverse photoautotrophic and photoheterotrophic conditions and perform nitrogen fixation.

### PHB production titer per liter is highest during photoheterotrophy and varies according to nitrogen source

We sought to understand PHB production during photoheterotrophy, in which cells use solar energy to break down organic substrates for carbon and electrons. We focused on sodium butyrate, a highly reduced organic compound known to be a preferred substrate for PHB production among related organisms (Ranaivoarisoa et al., 2019). Additionally, we grew the strains with two different nitrogen sources: N_2_ gas (i.e., nitrogen-fixing conditions) and ammonium chloride (NH_4_Cl). We generally observed the highest PHB titer per liter during photoheterotrophy, ranging from 4.96 ± 5.25 mg/L to 44.08 ± 1.77 mg/L (**Table 2, Figure 2**). Interestingly, nitrogen-fixing growth stimulated significantly more PHB production titer in both strains. *R. vannielii* produced 44.08 ± 1.77 mg/L during nitrogen fixation, compared to 8.25 ± 2.83 mg/L when supplied NH_4_Cl (*p* = 4.92E-05) (**Table 2, Figure 2**). Likewise, *R. udaipurense* produced 36.99 ± 6.97 mg/L during nitrogen fixation, compared to 4.86 ± 5.25 mg/L (*p* = 3.13E-03) (**Table 2, Figure 2**). Together, these data suggest that PHB production titer per liter of cell culture is highest during photoheterotrophy when cells are performing nitrogen fixation.

**Figure 2.**
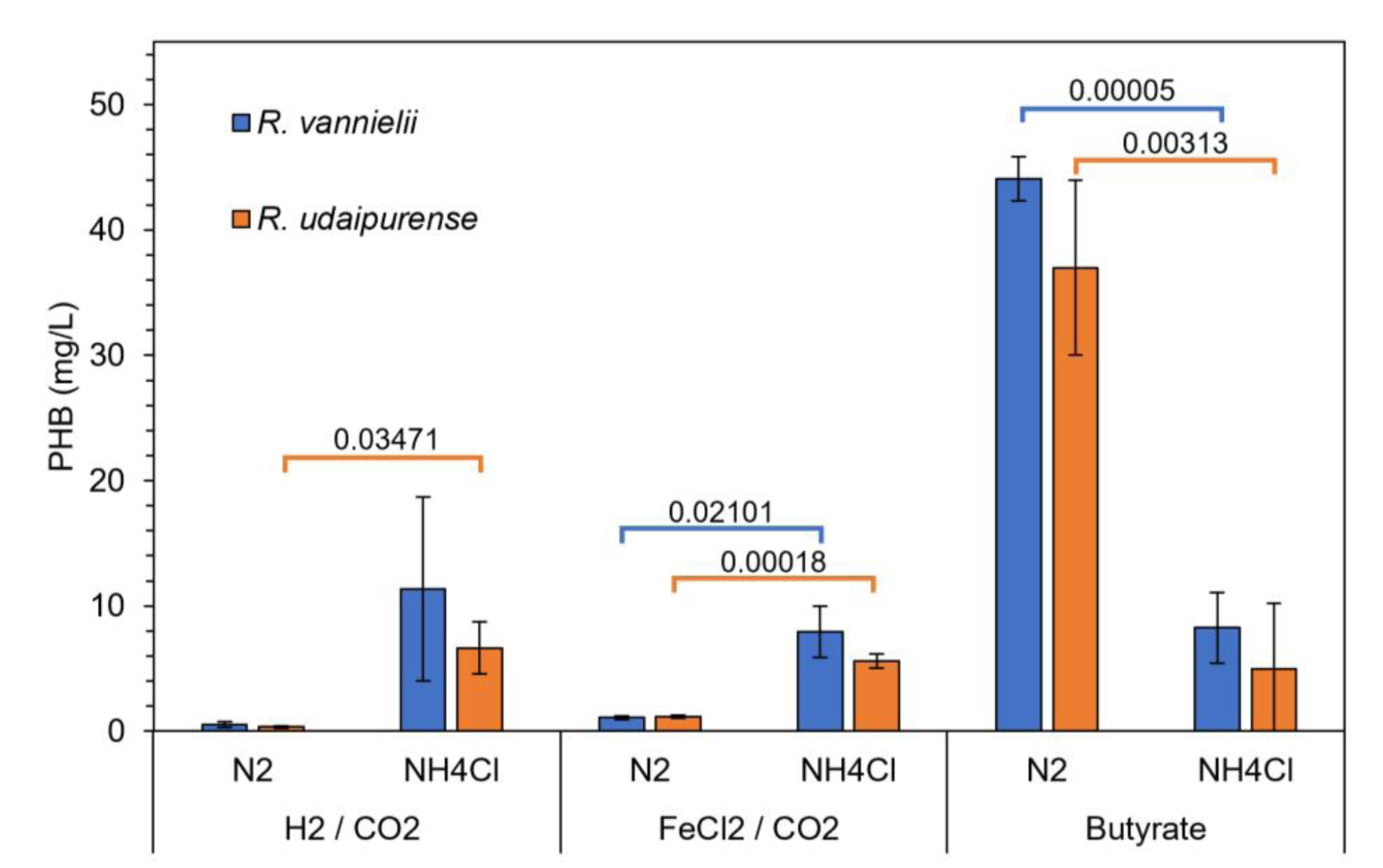
PHB production per liter by *Rhodomicrobium* spp. during photoautotrophic and photoheterotrophic growth. Intracellular PHB production (mg of PHB per liter of cell culture) during photoautotrophic growth with the following electron and carbon sources: hydrogen (H_2_) gas and carbon dioxide (CO_2_) (i.e., photoautotrophy), ferrous chloride (FeCl_2_) and CO_2_ (photoferrotrophy), and sodium butyrate (photoheterotrophy). **N_2_**: nitrogen-fixing growth with dinitrogen gas as sole nitrogen source. **NH_4_Cl:** non-nitrogen fixing growth with ammonium chloride as sole nitrogen source. *P-*values are calculated between N_2_ and NH_4_Cl growth conditions. Significantly different groups are denoted with horizontal bars and *p-*values.

**Table 2.** PHB production, carbon conversion, and electron conversion for *R. vannielii* and *R. udaipurense* under different metabolic regimes

| Strain | Electron Donor/Substrate | Nitrogen Source | PHB (mg/L) | *P*-value | PHB (mg/L/cell) * 10E-7 | *P*-value | Specific productivity (PHB/mg/L/cell/h) * 10E-9 | *P*-value | PHB carbon yield (%) | *P*-value (N<sub>2</sub> v. NH<sub>4</sub>) | PHB electron yield (%) | *P*-value |
| --- | --- | --- | --- | --- | --- | --- | --- | --- | --- | --- | --- | --- |
| *R. vannielii* | H<sub>2</sub> / CO<sub>2</sub> | N<sub>2</sub> | 0.53 (0.23) | 0.06 | 0.59 (0.28) | 0.08 | 0.25 (0.12) | 0.08 | 0.34 (0.16) | 0.08 | 6.72 (2.91) | 0.06 |
|  |  | NH<sub>4</sub>Cl | 11.35 (7.33) |  | 9.67 (6.8) |  | 2.92 (2.06) |  | 27.56 (21.06) |  | 86.57 (55.96) |  |
|  | FeCl<sub>2</sub> / CO<sub>2</sub> | N<sub>2</sub> | 1.09 (0.13) | **0.02** | 1.6 (0.18) | **0.02** | 0.37 (0.04) | **0.02** | 0.69 (0.17) | **0.02** | 13.98 (2.42) | **0.05** |
|  |  | NH<sub>4</sub>Cl | 7.92 (2.04) |  | 16.24 (3.81) |  | 3.11 (0.73) |  | 6.03 (1.38) |  | 35.88 (9.2) |  |
|  | Butyrate | N<sub>2</sub> | 44.08 (1.77) | **4.92E-05** | 20.91 (0.13) | **7.07E-04** | 12.6 (0.08) | 0.31 | 9.67 (0.87) | **1.12E-04** | 1.17 (0.05) | **2.60E-05** |
|  |  | NH<sub>4</sub>Cl | 8.25 (2.83) |  | 7.27 (2.5) |  | 10.23 (3.53) |  | 1.42 (0.37) |  | 0.21 (0.06) |  |
|  | Carbon Felt / CO<sub>2</sub> | N<sub>2</sub> | 0.13 (0.03) | 0.13 | 0.39 (0.14) | **0.05** | 0.23 (0.08) | **0.05** | 0.11 (0.01) | 0.90 | TBD | TBD |
|  |  | NH<sub>4</sub>Cl | 0.08 (0.03) |  | 0.11 (0.05) |  | 0.07 (0.03) |  | 0.12 (0.08) |  | 6.4 (2.09) |  |
| *R. udaipurensis* | H<sub>2</sub> / CO<sub>2</sub> | N<sub>2</sub> | 0.35 (0.09) | **0.03** | 0.47 (0.09) | 0.07 | 0.2 (0.04) | 0.08 | 0.24 (0.07) | **0.04** | 4.46 (1.19) | **0.04** |
|  |  | NH<sub>4</sub>Cl | 6.65 (2.1) |  | 6.95 (3.2) |  | 2.1 (0.97) |  | 19.75 (7.27) |  | 50.78 (15.99) |  |
|  | FeCl<sub>2</sub> / CO<sub>2</sub> | N<sub>2</sub> | 1.16 (0.14) | **1.82E-04** | 2.16 (0.25) | **9.38E-04** | 0.41 (0.05) | **4.19E-02** | 0.68 (0.04) | **1.71E-04** | 18.87 (6.62) | 0.07 |
|  |  | NH<sub>4</sub>Cl | 5.61 (0.56) |  | 5.03 (0.51) |  | 1.23 (0.47) |  | 4.25 (0.45) |  | 28.62 (1.84) |  |
|  | Butyrate | N<sub>2</sub> | 36.99 (6.97) | **3.13E-03** | 18.45 (1.68) | **0.02** | 15.25 (1.39) | **3.17E-02** | 6.29 (1.24) | **3.55E-03** | 0.75 (0.14) | **4.26E-03** |
|  |  | NH<sub>4</sub>Cl | 4.96 (5.25) |  | 5.08 (5.52) |  | 7.16 (7.77) |  | 0.91 (0.87) |  | 0.13 (0.12) |  |
|  | Carbon Felt / CO<sub>2</sub> | N<sub>2</sub> | 0.08 (0.02) | **0.04** | 0.16 (0.04) | **0.03** | 0.1 (0.02) | **0.03** | 0.04 (0.02) | 0.20 | TBD | TBD |
|  |  | NH<sub>4</sub>Cl | 0.04 (0.01) |  | 0.08 (0.02) |  | 0.05 (0.01) |  | 0.07 (0.02) |  | 6.91 (1.64) |  |

N<sub>2</sub>: nitrogen-fixing growth with dinitrogen gas as sole nitrogen source. NH<sub>4</sub>Cl: non-nitrogen fixing growth with ammonium chloride as sole nitrogen source. *P*-values are calculated between N<sub>2</sub> and NH<sub>4</sub>Cl growth conditions. Significant *P*-values (<0.05) are bolded. () = standard error values from n ≥ 3.

We then explored PHB production during photoautotrophic growth, wherein cells use solar energy to drive CO_2_ fixation. Here, we tested two distinct electron donors, hydrogen gas (H_2_) and ferrous iron (Fe_2+_, supplied as FeCl_2_ along with nitrilotriacetic acid, an iron chelating agent). Altogether, we observed PHB production titer ranging from 0.35 ± 0.09 mg/L to 11.35 ± 7.33 mg/L across all photoautotrophic growth conditions tested (**Table 2, Figure 2**). Nitrogen fixation generally resulted in less PHB production titer per liter: we observed PHB production titer ranging from 0.35 ± 0.09 mg/L to 1.16 ± 0.14 mg/L when cultures were supplied N_2_ gas as a nitrogen source. Conversely, cultures supplemented with NH_4_Cl produced between 5.61 ± 0.56 mg/L and 11.35 ± 7.33 mg/L (**Table 2, Figure 2**). This suggests that photoautotrophic PHB production titer per liter of cell culture is greatest during non-nitrogen fixing growth.

Finally, we examined PHB accumulation during photoelectroautrophic growth. Here, cells conducting photosynthetic CO_2_ fixation use a poised electrode as an electron source. PHB production titer on a per liter basis was low, ranging from 0.04 (± 0.01) mg/L to 0.13 (± 0.03) mg/L (**Table 2, Figure 3**).

**Figure 3.**
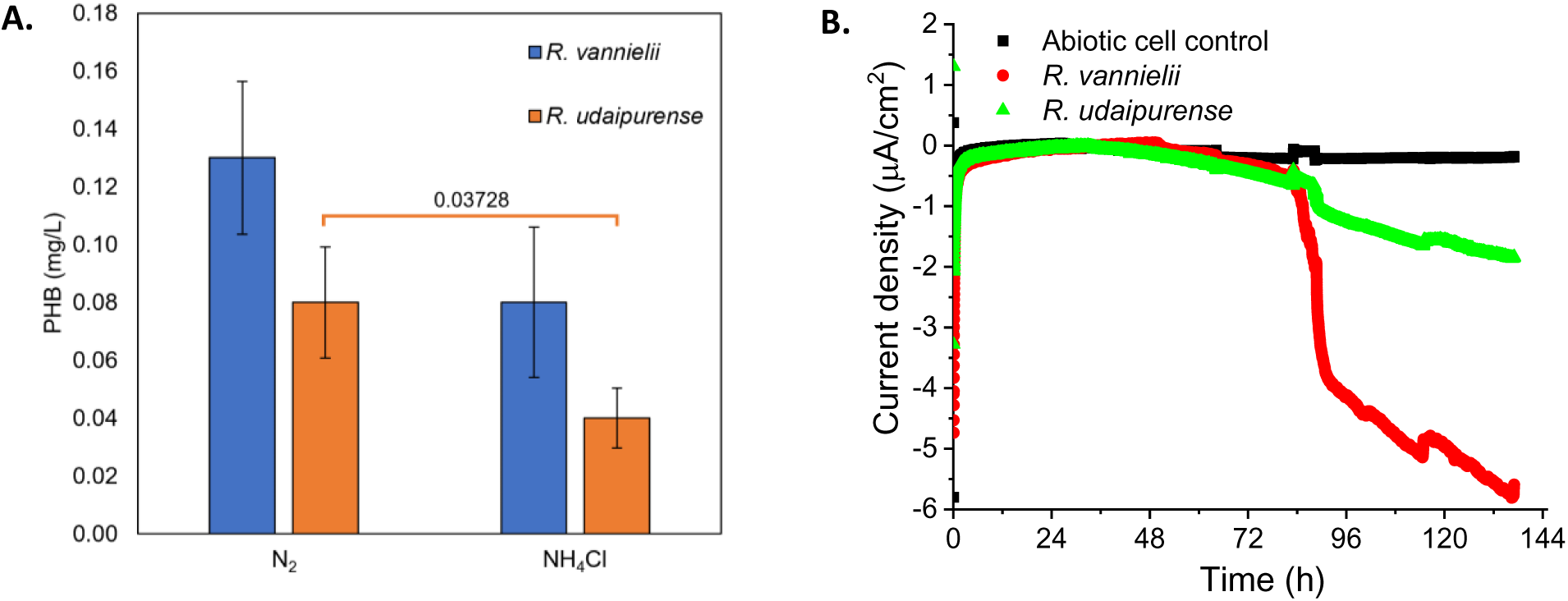
PHB production per liter by *Rhodomicrobium* spp. during photoelectrotrophy. **A.** Intracellular PHB production (mg of PHB per liter of cell culture) during photoelectrotrophic growth using a carbon felt cathode poised at +100 mV vs. standard hydrogen electrode. **N_2_**: nitrogen-fixing growth with dinitrogen gas as sole nitrogen source. **NH_4_Cl:** non-nitrogen fixing growth with ammonium chloride as sole nitrogen source. *P-*values are calculated between N_2_ and NH_4_Cl growth conditions. Significantly different groups are denoted with horizontal bars and *p-*values. **B.** Current uptake by *R. vannielii* and *R. udaipurense* in microbial electrosynthesis reactors supplemented with NH_4_Cl.

### Normalized PHB production and specific productivity trends align with titer

Because cell growth dynamics vary across culture conditions, we normalized PHB production according to cell number. Photoheterotrophy showed some of the highest PHB production per liter per cell, ranging from 5.08 * 10E-07 mg/L/cell to 20.91 * 10E-07 mg/L/cell (**Table 2**).

We also determined specific productivity, defined as PHB mg/L/cell/h. As with other metrics, we find that photoheterotrophic growth stimulates the highest specific productivity (**Table 2**). Taken together, these normalized metrics suggest that photoheterotrophic growth on butyrate generally stimulates the most PHB production. However, some overlap among the photoheterotrophic and photoautotrophic growth conditions suggests that the amount of PHB produced is not strictly a function of carbon and electron sources, and that other factors may be at play.

As with PHB production titer per liter, we assessed the influence of nitrogen fixation on PHB production. We generally observed an influence of nitrogen source on PHB production across all three productivity metrics. While these differences were not always statistically significant between both strains, we did observe a few consistent trends. First, photoheterotrophic growth on N_2_ gas resulted in significantly more PHB production per cell compared to non-fixing growth (*R. vannielii, p* = 7.07E-04; *R. udaipurense, p* = 0.02) (**Table 2, Figure 2**). In fact, photoheterotrophic growth on N_2_ gas showed the highest PHB production compared to all other conditions across all three productivity metrics (**Table 2**) We observed the inverse trend during photoferrotrophic growth, with a statistically significant increase in normalized PHB production and specific productivity in the non-fixing condition compared to growth on N_2_ (**Table 2**). Altogether, these data suggest that nitrogen source influenced PHB accumulation, but that this role varies according to other metabolic factors like carbon and electron source.

### *Rhodomicrobium* spp. produce low PHB titers via microbial electrosynthesis

Microbial electrosynthesis (MES) is a method of producing bioproducts by taking advantage of bacteria that accept electrons from poised cathodes. Here, we constructed an anoxic MES system with a carbon felt cathode poised at +100 mV continuously for 144 hours. We supplied energy via light and carbon via CO_2_. At the end of the experiment, all cells comprising the planktonic and biofilm populations were assessed for PHB production, and total current uptake over the duration of the experiment was calculated. We observed enhanced current uptake in all biological reactors relevant to abiotic controls, with a maximum current density of approximately −6 µA/cm^2^ for *R. vannielii* and −2 µA/cm^2^ for *R. udaipurense* under non-nitrogen fixing conditions (**Figure 3B**). Total current uptake reached approximately −2 Coulombs for *R. vannielii* and −1 Coulombs for *R. udaipurense* under non-fixing conditions (**Supplemental Figure S1**). We measured PHB production titer from 0.04 ± 0.01 mg/L to 0.13 ± 0.03 mg/L between both organisms and nitrogen regimes (**Table 2, Figure 3A**). While we observed greater PHB accumulation in both strains during N_2_ fixation, *R. udaipurense* showed a significant difference in PHB production titer per liter between N_2_ fixing and non-fixing growth during photoelectrotrophy (*p* = 0.04) (**Table 2, Figure 3A**).

We also normalized PHB production to cell number. Compared to other conditions, photoelectrotrophic growth showed the lowest normalized PHB production, ranging from 0.08 ± 0.02 mg/L to 0.39 ± 0.14 mg/L (**Table 2**). Only *R. udaipurense* demonstrated a statistically significant influence of nitrogen source on normalized productivity, with N_2_ fixing growth leading to 0.16 ± 0.04 mg/L compared to 0.08 ± 0.02 mg/L for non-fixing growth. Finally, we calculated specific productivity (mg/L/cell/h) and found similarly low values ranging from 0.05 ± 0.02 mg/L to 0.23 ± 0.08 mg/L with *R. udaipurense* showing significantly higher specific productivity during N_2_ fixing growth (*p* = 0.03) (**Table 2**). Compared to other growth conditions across all productivity metrics, photoelectrotrophic growth is the least productive condition. Furthermore, N_2_-fixing growth may lead to enhanced production, though the results here are mixed.

### Carbon and electron yields are not correlated with overall PHB production

We next sought to understand the efficiency with which cells convert carbon and electrons into intracellular PHB. To assess carbon conversion, we first determined the initial and final amounts of carbon mol in the system (both organic and inorganic). We then calculated the total carbon mol in the observed PHB. The ratio between the consumed carbon mol and observed carbon mol in PHB defines the overall percent carbon yield (see **Supplemental Methods for detailed calculations**). Following these calculations, we find the highest carbon yields of 27.56% ± 21.06% (R*. vannielii*) and 19.75% ± 7.27% (*R. udaipurense*) during photoautotrophic growth with H_2_ and NH_4_Cl, while the lowest carbon yields are among the electrotrophic growth conditions (0.04% ± 0.02% to 0.12% ± 0.08%) (**Table 2**). Photoheterotrophic growth on butyrate showed a range of carbon yields; when grown in N_2_-fixing conditions, *R. vannielii* and *R. udaipurense* showed carbon yields of 9.67% ± 0.87% and 6.29% ± 1.24%, whereas non-fixing growth showed yields of only 1.42% ± 0.37% and 0.91 ± 0.87%, respectively.

To determine whether either strain shows a statistically significant difference in carbon yields across our tested growth conditions, we performed a one-way ANOVA. We observed a statistically significant difference in the group means of *R. vannielii* (F(7,17) = 4.25, *p* = 0.007) and *R. udaipurense* (F(7,16) = 19.79, *p =* 9.23E-07) (**Supplemental Tables 1 and 2**). Games-Howell post hoc analysis revealed significantly greater carbon yields during N_2_-fixing photoheterotrophy in *R. vannielii* compared to most other growth conditions, apart from FeCl_2_ + NH_4_Cl and H_2_ + NH_4_Cl, which showed high variability in carbon yields (**Supplemental Table 3**). We also observed statistically significant differences in the *R. udaipurense* carbon yields during photoferrotrophy compared to H_2_ + N_2_ and both photoelectrotrophic conditions, which had the lowest carbon yields (**Table 2, Supplemental Table 4**). Altogether, these data suggest that *Rhodomicrobium* strains generally divert a small fraction of carbon toward PHB accumulation; all growth conditions apart from H_2_ + NH_4_Cl showed carbon yields <10% (**Table 2**). Importantly, our data suggests that more carbon goes toward PHB accumulation under certain photoautotrophic (H_2_ + NH_4_Cl) and photoheterotrophic (Butyrate + N_2_) conditions.

We took a similar approach to calculating electron yield based on the ratio of consumed electrons to the electrons predicted to be directed toward PHB production (see **Supplemental Methods** for detailed calculations). Both strains showed the highest electron yield among photoautotrophic growth conditions, with the lowest electron yields observed during photoheterotrophic growth (**Table 2**). Among all conditions, growth on H_2_ + NH_4_Cl showed the highest electron yields of 86.57% ± 55.96% and 50.78% ± 15.99% for *R. vannielii* and *R. udaipurense,* respectively (**Table 2**). Photoheterotrophic electron yields ranged from 0.13% ± 0.12% (*R. vannielii* + Butyrate + NH_4_Cl) to 1.17% ± 0.05% (*R. vannielii +* Butyrate + N_2_) (**Table 2**). One-way ANOVA confirmed that there are significant differences among the respect electron yields of *R. vannielii* (F(7,15) = 2.71, *p* = 0.005) and *R. udaipurense* (F(6,14) = 2.85, *p =* 1.43E-06) (**Supplemental Tables 5 and 6**). Games-Howell post hoc testing suggests that electron yields are statistically similar across most growth conditions, though photoferrotrophy appears to stimulate a statistically greater electron yield in *R. udaipurense* compared to photoheterotrophy and photoelectrotrophy (**Supplemental Tables 7 and 8**). As with carbon yield, substantial variability in the photoautotrophic growth conditions with H_2_ and NH_4_Cl make it difficult to draw conclusions about the apparently high electron yield in that condition. Despite this, it appears that a greater fraction of available electrons is diverted toward PHB during photoautotrophy compared to photoheterotrophy.

Taken together, the carbon and electron yield data imply a complex dynamic between carbon and electron conversion in the context of PHB accumulation, with photoheterotrophy favoring broadly favoring carbon yield and photoautotrophy favoring electron yield. To better understand the relationship between carbon yield, electron yield, and PHB accumulation, we conducted linear regression analyses. Neither carbon yield (R_2_ = 0.22, *p* = 0.07) nor electron yield (R_2_ = 0.01, *p* = 0.70) showed a significant correlation with PHB productivity (mg/L/cell) (**Figure 4, Supplemental Tables 9 and 10**).

**Figure 4.**
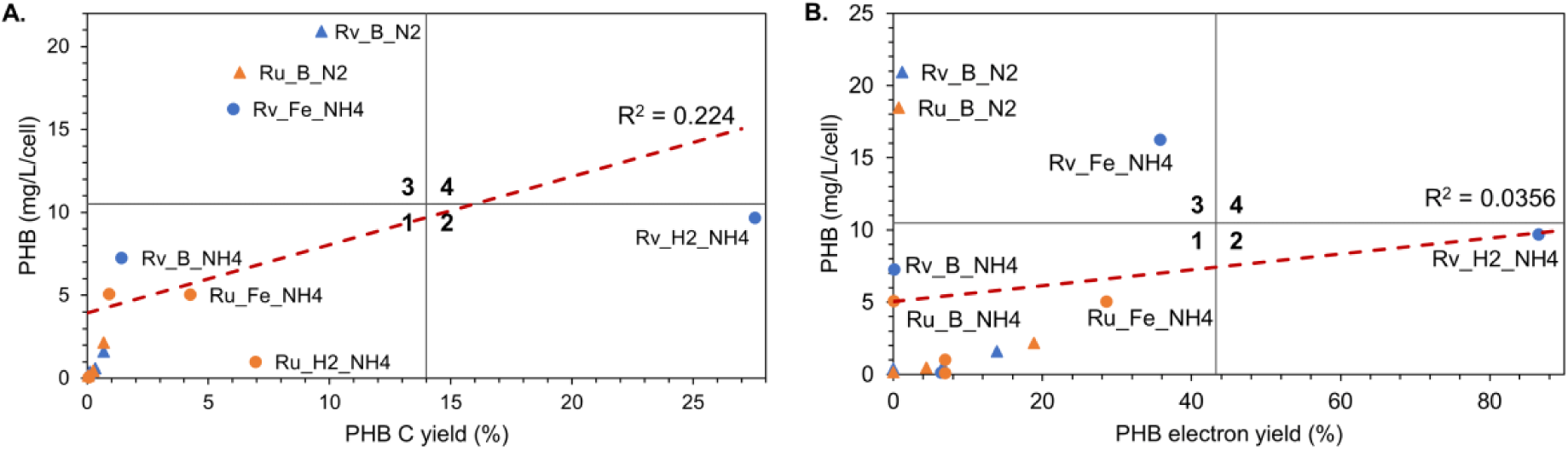
Carbon and electron yields compared to normalized PHB production. **A.** PHB carbon yield (%) and **B.** PHB electron yield (%) vs. normalized PHB production. Samples have been grouped according to four classes by bisecting the horizonal and vertical axes. Trendline and corresponding R_2_ value reflects all samples. Blue, Rv: *R. vannielii*. Orange, Ru: *R. udaipurense*. B: butyrate. H2: hydrogen electron donor. Fe: FeCl_2_ electron donor. Triangles, N2: nitrogen gas. Circles, NH4: ammonium chloride.

We then plotted the carbon and electron yield values against the normalized PHB production, allowing us to categorize each sample according to their carbon/electron conversion and PHB production dynamics. We chose to distinguish between groups with high and low yields by bisecting each axis at one-half of the largest value in either direction. This resulted in four classes of PHB producer: 1) low carbon/electron yield and low PHB yield, 2) high carbon/electron yield and low PHB yield, 3) low carbon/electron yield and high PHB yield, and 4) high carbon/electron yield and high PHB production. According to this framing, Class 4 producers are ideal, as these produce relatively large amounts of PHB with greater carbon or electron conversion efficiency. Conversely, Class 1 may be less preferred due to low production and conversion efficiencies. Here, we find that none of our conditions fall into Class 4. However, N_2_-fixing photoheterotrophy shows promise as a Class 3 producer with high PHB production with relatively moderate carbon/electron yields, while non-fixing photoautotrophy shows promise with higher carbon and electron yields than other conditions. It is important to note that *R. vannielii* grown in non-fixing autotrophic conditions (i.e., H_2_ + NH_4_Cl) likely skews these data, as it has comparatively high carbon and electron yields with substantial variability (27.56% ± 21.06% carbon yield, 86.57% ± 55.96% electron yield) (**Table 2**).

## Discussion

Conventional plastics impose enormous costs throughout their life cycle. Novel polymers like PHB can upend this cycle if they fulfill certain requirements, chief among them being sustainable, economically viable production via renewable inputs. Raw materials account for over half of production costs for biopolymers (Reshmy et al., 2021); 70-80% of those costs derive from the carbon source alone (Sirohi et al., 2021). Here, we show that PHB can be produced using diverse carbon sources with minimal nutrients and energy from light. Unsurprisingly, we observed the greatest overall PHB production (up to 44.08 mg/L) when an organic substrate (butyrate) was provided. While arguably the most expensive carbon substrate we examined, butyrate has been shown to be an ideal substrate for PHB production among related photosynthetic purple non-sulfur bacteria (Carlozzi et al., 2019; Luongo et al., 2017; Ranaivoarisoa et al., 2019). Furthermore, butyric acid can be sourced sustainably via microbial fermentation that uses biomass feedstocks like corn husk (Xiao et al., 2018), straw (Baroi et al., 2015), corn fiber (Zhu et al., 2002), oilseed rape straw (Huang et al., 2016), and sugarcane bagasse (Wei et al., 2013). Importantly, diverse biological chassis show varying preferences for substrates as it pertains to PHB production; many organic substrates like acetate, propionate, glutamate, and more, support PHB accumulation in phototrophs (Monroy and Buitrón, 2020). Therefore, heterotrophic PHB production remains promising, so long as substrates can be sourced cheaply and with minimal environmental impact.

The most sustainable carbon substrate – and the one that will have the greatest impact on climate change – is CO_2_. Here, we show that *Rhodomicrobium* produces PHB using only CO_2_ and light energy along with a variety of electron donors. While PHB titers were low (0.04 – 11.35 mg/L), we observed comparable carbon yields and superior electron yields with respect to photoheterotrophy. This strongly suggests photoautotrophy’s strength is in its sustainability and conversion efficiency. Therefore, photoautotrophic PHB production warrants additional exploration. One approach entails genetic modifications intended to enhance PHB titers. A recent study found that overexpressing the *Rubisco* form I or *Rubisco* form I and II genes enhances PHB production per cell almost twofold in *Rhodopseudomonas palustris* TIE-1 grown photoautotrophically with NH_4_Cl as a nitrogen source (Tahina Ranaivoarisoa 2023, unpublished results). The same study found that deleting the *nifA* gene responsible for encoding nitrogen-fixing machinery resulted in almost two-fold higher PHB production per cell when NH_4_Cl was provided. This suggests that enhancing photosynthetic CO_2_ fixation and/or deleting electron sinks like nitrogen fixation could enhance yields in other photosynthetic purple non-sulfur bacteria like *Rhodomicrobium*. Similar approaches should be explored in *Rhodomicrobium* spp., not only to design more efficient PHB producers, but to better understand the underlying physiology.

Additionally, our data lends credence to efforts exploring novel biological chassis for bioproduction. To the best of our knowledge, this is the first report of *Rhodomicrobium* strains producing PHB under diverse metabolic conditions. Previously, we explored PHB production in the photosynthetic purple non-sulfur bacteria *Rhodopseudomonas palustris* TIE-1 across the same growth conditions using identical analytical methods (Ranaivoarisoa et al., 2019). This methodological consistency allows us to directly compare the present data with those published by Ranaivoarisoa et al. (2019). In the present study, distantly related phototrophs from the *Rhodomicrobium* spp. produced comparable or superior PHB production titers, across all growth conditions. However, both *Rhodomicrobium* species demonstrated 5 to 6 orders of magnitude greater PHB per cell than TIE-1. This strongly suggests that, while the final titers were similar, *Rhodomicrobium* spp. accomplished this with far fewer cells, likely by accumulating more PHB per cell than TIE-1. One potential explanation may lie in electron yields. Half of our *Rhodomicrobium* cultures showed average electron yields greater than 6%, with some as high as 50 – 80%; conversely, the majority of TIE-1’s electron yields across varying growth conditions were below 1%, Interestingly, TIE-1 showed the highest electron yields during photoelectrotrophy, reaching 4.39% (NH_4_Cl) and 7.34% (N_2_), which is comparable to the electron yields observed in *Rhodomicrobium* (6.40% with NH_4_Cl, electron yield with N_2_ TBD). Superimposing the corresponding data from Ranaivoarisoa et al. (Ranaivoarisoa et al., 2019) onto our data in **Figure 4** reveals that TIE-1 falls in the Class I group for carbon and electron conversion across all comparable growth conditions (i.e., photoheterotrophy with butyrate, photoautotrophy with H_2_ gas, photoferrotrophy, and photoelectrotrophy, all with N_2_ or NH_4_Cl nitrogen sources) (**Figure 5**). This suggests that *Rhodomicrobium* may convert electrons into PHB more effectively than TIE-1 in certain contexts. Furthermore, it suggests that greater biomass would substantially increase PHB titers far beyond those observed in TIE-1.

**Figure 5.**
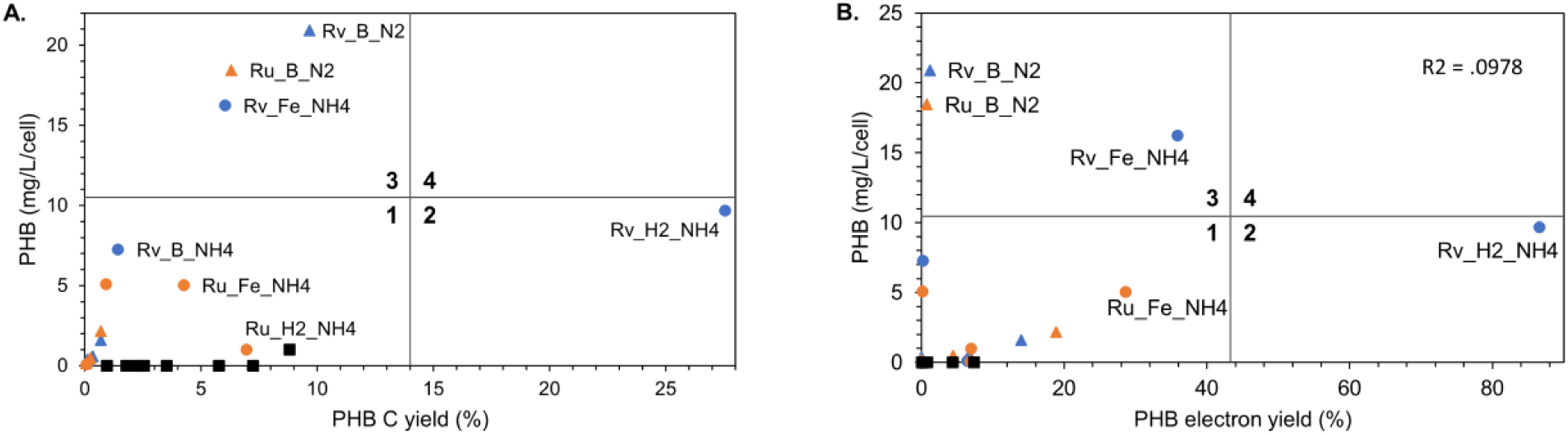
*Rhodomicrobium* spp. and TIE-1 carbon, and electron yields compared to normalized PHB production. **A.** PHB carbon yield (%) and **B.** PHB electron yield (%) vs. normalized PHB production. Samples have been grouped according to four classes by bisecting the horizonal and vertical axes. Black: *R. palustris* TIE-1 (*Ranaivoarisoa et al*., 2019). Blue, Rv: *R. vannielii*. Orange, Ru: *R. udaipurense*. B: butyrate. H2: hydrogen electron donor. Fe: FeCl_2_ electron donor. Triangles, N2: nitrogen gas. Circles, NH4: ammonium chloride

The comparison with TIE-1 is apt because it highlights the strengths and weaknesses of different non-model organisms. Since being isolated in 2005, *R. palustris* TIE-1 has been studied for its ability to conduct diverse metabolisms including extracellular electron uptake (EEU), iron oxidation, and photosynthetic carbon capture (Bose et al., 2014; Bose and Newman, 2011; Gupta et al., 2019; Jiao et al., 2005). These efforts have led to the development of genetic systems and protocols for growth and characterization. As such, TIE-1 is a reliable organism for investigating unique metabolisms like EEU and photoferrotrophy or bioproduction in purple non-sulfur bacteria. On the other hand, *Rhodomicrobium vannielii* and *Rhodomicrobium udaipurense* are severely understudied in this context. Consequently, reliable genetic systems are sparse, with few reports on successful genetic manipulation in this genus (Dziuba et al., 2023). It stands to reason that future efforts should focus on organisms like TIE-1 with well-established protocols. However, we argue that non-model organisms like *Rhodomicrobium* warrant further consideration in light of these findings. We report substantial discrepancies between the two organisms with respect to electron conversion and per cell yield, in addition to highly variable titers. Future studies should investigate the underlying causes of these differences, and this will require efforts to develop reliable genetic systems, high-resolution imaging protocols, and more tools in *Rhodomicrobium*. However, this does not discount the utility of an organism like TIE-1. In fact, TIE-1 presents an opportunity to probe fundamental questions about photosynthetic carbon capture, extracellular electron uptake, and photoferrotrophy with relative ease. These findings can inform studies in related non-model organisms with similar genetics and metabolisms, and vice versa. In this way, more established models like TIE-1 help to fine-tune experimentation in non-model organisms like *Rhodomicrobium*.

In summary, this study highlights two novel biological chassis for PHB production, *Rhodomicrobium vannielii* and *Rhodomicrobium udaipurense.* Future studies should explore methods for enhancing PHB production in these organisms, as well as identify other novel biological chassis with the ability to utilize inexpensive, sustainable inputs for bioplastic production.

## Acknowledgements

We would like to thank Tahina Ranaivoarisoa for reviewing mathematical equations and providing valuable feedback. This work was supported by the following grants to A.B.: The David and Lucile Packard Foundation Fellowship (201563111), the U.S. Department of Energy (grant number DESC0014613), and the U.S. Department of Defense, Army Research Office (grant number W911NF-18-1-0037), Gordon and Betty Moore Foundation, National Science Foundation (Grant Number 2021822, Grant Number 2124088, and Grant Number 2117198), the U.S. Department of Energy by Lawrence Livermore National Laboratory under Contract DEAC5207NA27344 (LLNL-JRNL-812309), an NIGMS grant (NIHR01GM141344), and a DEPSCoR grant (FA9550-21-1-0211). A.B. was also funded by a Collaboration Initiation Grant, an Office of the Vice-Chancellor of Research Grant, an International Center for Energy, Environment, and Sustainability Grant and a SPEED grant from Washington University in St. Louis. E.M.C. was supported by the Schneiderman Fellowship.

