## Supplemental Figure and Tables for "The phototrophic bacteria *Rhodomicrobium* spp. are novel chassis for bioplastic production"

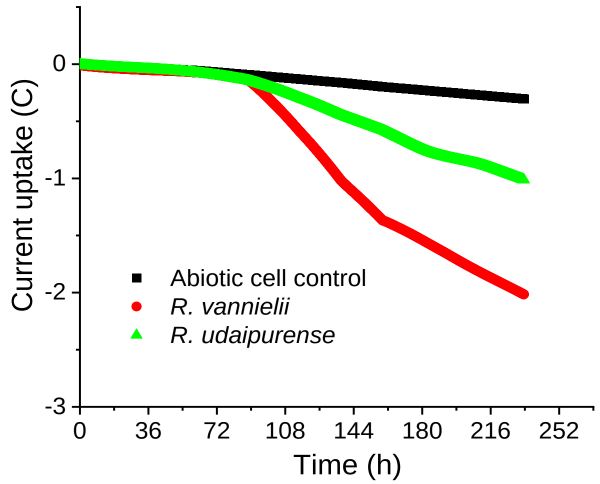


**Supplemental Figure S1.** Total current uptake (Coulombs) in microbial electrosynthesis reactors supplemented with $NH_4Cl$.

**Supplemental Table S1. One-way ANOVA of *R. vannielii* carbon yield**

| *Groups* | *Count* | *Sum* | *Average* | *Variance* |
|---|---|---|---|---|
| Rv H2N2 1 | 4 | 1.367924 | 0.341981 | 0.024485 |
| Rv FeN2 1 | 3 | 2.056502 | 0.685501 | 0.028067 |
| Rv BNH4 3 | 3 | 4.256128 | 1.418709 | 0.138957 |
| Rv BN2 3 | 3 | 29.01135 | 9.670451 | 0.755211 |
| Rv H2NH4 1 | 4 | 110.2225 | 27.55563 | 443.4161 |
| Rv ENH4 1 | 3 | 0.347872 | 0.115957 | 0.007193 |
| Rv FeNH4Cl 1 | 3 | 18.07937 | 6.026456 | 1.912628 |
| Rv EN2 1 | 2 | 0.214038 | 0.107019 | 9.1E-05 |

| *Source of Variation* | *SS* | *df* | *MS* | *F* | *P-value* | *F crit* |
|---|---|---|---|---|---|---|
| Between Groups | 2338.39 | 7 | 334.0557 | 4.25069 | 0.006968 | 2.614299 |
| Within Groups | 1336.006 | 17 | 78.58859 | | | |
| Total | 3674.396 | 24 | | | | |

**Supplemental Table S2. One-way ANOVA of *R. udaipurense* carbon yield**

| *Groups* | *Count* | *Sum* | *Average* | *Variance* | | |
|---|---|---|---|---|---|---|
| Ru H2N2 1 | 3 | 0.724513 | 0.241504 | 0.004767 | | |
| Ru FeN2 1 | 3 | 2.047337 | 0.682446 | 0.001753 | | |
| Ru BNH4 3 | 3 | 2.736771 | 0.912257 | 0.753553 | | |
| Ru BN2 3 | 3 | 18.88156 | 6.293855 | 1.546411 | | |
| Ru H2NH4 1 | 3 | 59.25541 | 19.7518 | 52.79594 | | |
| Ru ENH4 1 | 3 | 0.20482 | 0.068273 | 0.000417 | | |
| Ru FeNH4Cl 1 | 3 | 12.76292 | 4.254307 | 0.206484 | | |
| Ru EN2 1 | 3 | 0.134762 | 0.044921 | 0.00028 | | |
| *Source of Variation* | *SS* | *df* | *MS* | *F* | *P-value* | *F crit* |
| Between Groups | 957.6172 | 7 | 136.8025 | 19.78715 | 9.23E-07 | 2.657197 |
| Within Groups | 110.6192 | 16 | 6.9137 | | | |
| Total | 1068.236 | 23 | | | | |

**Supplemental Table S3. Games-Howell post hoc test on *R. vannielii* carbon yields.** Q values of significant comparisons where $Q > Q_{crit}$ at $\alpha = 0.05$. Statistically significant comparisons are bolded.

| | Rv H2N2 | Rv H2NH4 | Rv FeN2 | Rv FeNH4 | Rv BN2 | Rv BNH4 | Rv EN2 |
|---|---|---|---|---|---|---|---|
| **Rv H2NH4** | 3.655 | | | | | | |
| **Rv FeN2** | 3.905 | 3.609 | | | | | |
| **Rv FeNH4** | 10.02 | 2.883 | 9.391 | | | | |
| **Rv BN2** | **25.979** | 2.399 | **24.867** | 5.464 | | | |
| **Rv BNH4** | 6.649 | 3.509 | 4.394 | 7.879 | **21.375** | | |
| **Rv EN2** | 4.231 | 3.686 | 8.437 | 10.483 | **26.953** | 8.614 | |
| **Rv ENH4** | 3.463 | 3.685 | 7.429 | 10.448 | **26.803** | 8.347 | 0.255 |

**Supplemental Table S4. Games-Howell post hoc test on *R. udaipurense* carbon yields.** Q values of significant comparisons where $Q > Q_{crit}$ at $\alpha = 0.05$. Statistically significant comparisons are bolded.

| | **Ru H2N2** | **Ru H2NH4** | **Ru FeN2** | **Ru FeNH4** | **Ru BN2** | **Ru BNH4** | **Ru EN2** |
|---|---|---|---|---|---|---|---|
| **Ru H2NH4** | 6.576 | | | | | | |
| **Ru FeN2** | **13.376** | 6.428 | | | | | |
| **Ru FeNH4** | **21.385** | 5.214 | **19.173** | | | | |
| **Ru BN2** | 11.903 | 4.471 | 11.046 | 3.773 | | | |
| **Ru BNH4** | 1.886 | 6.306 | 0.647 | 8.354 | 8.692 | | |
| **Ru EN2** | 6.777 | 6.643 | **34.628** | **22.675** | 12.307 | 2.446 | |
| **Ru ENH4** | 5.893 | 6.635 | **32.29** | **22.542** | 12.261 | 2.38 | 2.165 |

**Supplemental Table S5. One-way ANOVA of *R. vannielii* electron yield**

| *Groups* | *Count* | *Sum* | *Average* | *Variance* | | |
|---|---|---|---|---|---|---|
| Rv H2N2 1 | 4 | 26.87431 | 6.718576 | 8.472538 | | |
| Rv FeN2 1 | 3 | 41.92999 | 13.97666 | 5.86902 | | |
| Rv BNH4 3 | 3 | 0.63651 | 0.21217 | 0.003585 | | |
| Rv BN2 3 | 3 | 3.514651 | 1.17155 | 0.002205 | | |
| Rv H2NH4 1 | 4 | 346.2965 | 86.57412 | 3131.483 | | |
| Rv ENH4 1 | 3 | 19.20107 | 6.400357 | 4.357631 | | |
| Rv FeNH4Cl 1 | 3 | 107.6296 | 35.87653 | 84.6414 | | |
| Rv EN2 1 | 0 | 0 | TBD | TBD | | |
| *Source of Variation* | *SS* | *df* | *MS* | *F* | *P-value* | *F crit* |
| Between Groups | 21769.95 | 7 | 3109.993 | 4.854501 | 0.004966 | 2.706627 |
| Within Groups | 9609.615 | 15 | 640.641 | | | |
| Total | 31379.56 | 22 | | | | |

**Supplemental Table S6. One-way ANOVA of *R. udaipurense* electron yield**

| *Groups* | *Count* | *Sum* | *Average* | *Variance* | | |
|---|---|---|---|---|---|---|
| Ru H2N2 1 | 3 | 13.37832 | 4.459441 | 1.408011 | | |
| Ru FeN2 1 | 3 | 56.59857 | 18.86619 | 43.82257 | | |
| Ru BNH4 3 | 3 | 0.383398 | 0.127799 | 0.013251 | | |
| Ru BN2 3 | 3 | 2.240354 | 0.746785 | 0.020359 | | |
| Ru H2NH4 1 | 3 | 152.3461 | 50.78202 | 255.629 | | |
| Ru ENH4 1 | 3 | 20.73273 | 6.910909 | 2.679728 | | |
| Ru FeNH4Cl 1 | 3 | 85.86629 | 28.6221 | 3.387585 | | |
| *Source of Variation* | *SS* | *df* | *MS* | *F* | *P-value* | *F crit* |
| Between Groups | 6232.17 | 6 | 1038.695 | 23.68665 | 1.43E-06 | 2.847726 |
| Within Groups | 613.921 | 14 | 43.8515 | | | |
| Total | 6846.091 | 20 | | | | |

**Supplemental Table S7. Games-Howell post hoc test on *R. vannielii* carbon yields.** Q values of significant comparisons where $Q > Q_{crit}$ at $\alpha = 0.05$. Statistically significant comparisons are bolded.

| | Rv H2N2 | Rv H2NH4 | Rv FeN2 | Rv FeNH4 | Rv BN2 | Rv BNH4 | Rv EN2 |
|---|---|---|---|---|---|---|---|
| **Rv H2NH4** | 4.03 | | | | | | |
| **Rv FeN2** | 5.085 | 3.664 | | | | | |
| **Rv FeNH4** | 7.487 | 2.517 | 5.638 | | | | |
| **Rv BN2** | 5.389 | 4.316 | 12.944 | 9.239 | | | |
| **Rv BNH4** | 6.32 | 4.365 | **13.912** | 9.495 | **30.882** | | |
| **Rv EN2** | TBD | TBD | TBD | TBD | TBD | TBD | |
| **Rv ENH4** | 0.238 | 4.048 | 5.803 | 7.653 | 6.133 | 7.258 | TBD |

**Supplemental Table S8. Games-Howell post hoc test on *R. udaipurense* carbon yields.** Q values of significant comparisons where $Q > Q_{crit}$ at $\alpha = 0.05$. Statistically significant comparisons are bolded.

| | **Ru H2N2** | **Ru H2NH4** | **Ru FeN2** | **Ru FeNH4** | **Ru BN2** | **Ru BNH4** | **Ru EN2** |
|---|---|---|---|---|---|---|---|
| **Ru H2NH4** | 7.077 | | | | | | |
| **Ru FeN2** | 5.247 | 4.517 | | | | | |
| **Ru FeNH4** | **27.027** | 3.372 | 3.477 | | | | |
| **Ru BN2** | 7.609 | 7.665 | 6.703 | **36.986** | | | |
| **Ru BNH4** | 8.9 | 7.76 | 6.932 | **37.847** | 8.27 | | |
| **Ru EN2** | TBD | TBD | TBD | TBD | TBD | TBD | |
| **Ru ENH4** | 2.97 | 6.686 | 4.294 | **21.59** | 9.188 | 10.124 | TBD |

**Supplemental Table S9. Linear regression of PHB carbon yield (%) vs. normalized PHB (mg/L/cell * 10E-7)**

| ***Regression Statistics*** | |
|---|---|
| Multiple R | 0.471636289 |
| R Square | 0.22244079 |
| Adjusted R Square | 0.166900846 |
| Standard Error | 6.364165773 |
| Observations | 16 |

| **ANOVA** | *df* | *SS* | *MS* | *F* | *Significance F* |
|---|---|---|---|---|---|
| Regression | 1 | 162.2153548 | 162.2153548 | 4.005059695 | 0.065137612 |
| Residual | 14 | 567.0364839 | 40.50260599 | | |
| Total | 15 | 729.2518387 | | | |

| | *Coefficients* | *Standard Error* | *t Stat* | *P-value* | *Lower 95%* | *Upper 95%* | *Lower 95.0%* | *Upper 95.0%* |
|---|---|---|---|---|---|---|---|---|
| Intercept | 3.938398126 | 1.881350029 | 2.093389356 | 0.055001412 | -0.096696372 | 7.973492624 | -0.096696372 | 7.973492624 |
| PHB carbon yield (%) | 0.411265888 | 0.205503012 | 2.001264524 | 0.065137612 | -0.029494237 | 0.852026014 | -0.029494237 | 0.852026014 |

**Supplemental table S10. Linear regression of PHB electron yield (%) vs. normalized PHB (mg/L/cell * 10E-07)**

| *Regression Statistics* | |
|---|---|
| Multiple R | 0.113963393 |
| R Square | 0.012987655 |
| Adjusted R Square | -0.069263374 |
| Standard Error | 7.343904211 |
| Observations | 14 |

| **ANOVA** | *df* | *SS* | *MS* | *F* | *Significance F* |
|---|---|---|---|---|---|
| Regression | 1 | 8.51615208 | 8.51615208 | 0.157902644 | 0.698072292 |
| Residual | 12 | 647.1951487 | 53.93292906 | | |
| Total | 13 | 655.7113008 | | | |

| | *Coefficients* | *Standard Error* | *t Stat* | *P-value* | *Lower 95%* | *Upper 95%* | *Lower 95.0%* | *Upper 95.0%* |
|---|---|---|---|---|---|---|---|---|
| Intercept | 6.148739439 | 2.490444174 | 2.468932853 | 0.029551185 | 0.722527723 | 11.57495116 | 0.722527723 | 11.57495116 |
| PHB electron yield (%) | 0.032618526 | 0.082086102 | 0.397369656 | 0.698072292 | -0.146231726 | 0.211468779 | -0.146231726 | 0.211468779 |

**Supplemental Table S11. Theoretical total available electron(s) mol/mol substrate**

| Electron source | Total available electron(s)mol/Mol substrate |
|---|---|
| Succinate | 14 |
| Butyrate | 20 |
| Hydroxybutyrate | 18 |
| $Fe^{2+}$ | 1 |
| $H_2$ | 2 |

**Supplemental Table S12. Oxidation/Reduction values and theoretical electrons required for crotonic acid synthesis**

| Electron source | Oxidation/reduction value | Electrons required for crotonic acid synthesis |
|---|---|---|
| Succinate | +1 | 4 |
| Butyrate | -1 | 2 |
| Hydroxybutyrate | -0.5 | Not thermodynamically calculable |
| $CO_2$ | +4 | 18 |
| Crotonic acid | -0.5 | Not applicable |
