## Supplemental Methods for "The phototrophic bacteria *Rhodomicrobium* spp. are novel chassis for bioplastic production"

### PHB extraction and analysis

PHB extraction was performed as described previously (Ranaivoarisoa et al., 2019). Briefly, samples were dried for 6 hours under vacuum in a Savant SC210A Speedvac concentrator (Thermo Fisher Scientific Inc, USA). 425 µL of methanol, 500 µL of HPLC grade chloroform and 75 µL of 95-98% sulfuric acid were added to the dried samples. Samples were digested for 1 hour in a water bath at 95°C. Digested samples were cooled on ice rapidly. 0.5 mL of LC-MS grade water was then added. Samples were vortexed and centrifuged at 5000 x *g* for 10 min. The organic phases were transferred into glass vials and dried for 30 min via speed vacuum. Finally, the dried samples were re-suspended in 500 µL of 50% acetonitrile + 50% water. Samples were filtered with 0.22 µm PTFE membrane filter to remove any cell debris prior to analysis. PHB measurements were performed using LC-MS.

The % carbon yield was calculated using Equations 1, 2 and 3 below:

$$C\ mol\ substrate = \frac{Consumed\ substrate\ \left(\frac{g}{L}\right) \times Number\ of\ Carbon\ in\ substrate}{MW\ substrate} \quad (1)$$

$$C\ mol\ PHB = \frac{PHB\ as\ crotonic\ acid\ \left(\frac{g}{L}\right) \times Number\ of\ Carbon\ in\ crotonic\ acid}{MW\ crotonic\ acid} \quad (2)$$

$$\%\ C\ yield = 100 \times \frac{\text{C mol PHB}}{C\ mol\ substrate} \quad (3)$$

The % electron yield was calculated as described previously (Bai et al., 2021). The total available electrons obtained from oxidation of each organic acid to $CO_2$ are shown in **Supplemental Table S11**. The inorganic electron donor $Fe^{2+}$ and $H_2$ release 1 and 2 electrons respectively as shown in Supplemental Table S3 adapted from (Nevin et al., 2011). Electrons supplied for photoelectrotrophic growth were calculated directly from bioelectrochemical experiments wherein total current uptake was integrated over the operational time. Total electron uptake was used to calculate the % electron yield because the electrode is the direct electron donor under this growth condition; this prevents us from using the oxidation/reduction method described below for % electron yield calculations.  Electrons required for crotonic acid production were calculated from the oxidation/reduction value of the carbon in the conversion reaction of each substrate to crotonic acid (**Supplemental Table S12**). For example: in the conversion of succinate into crotonic acid, the oxidation/reduction value of butyrate is -1 and for crotonic acid it is -0.5, hence 2 mol $e^-$ are involved. To obtain the number of electrons (mol) in the consumed substrate, the number of mol of consumed substrate was multiplied by the theoretical total available electron when the substrate is fully oxidized to $CO_2$ (Equation 4). To obtain the number of electrons required for PHB production, the measured mol of crotonic acid was multiplied by the theoretical number of electrons required for one mol of crotonic acid (Equation 5). The obtained number was divided by the number of mol $e^-$ consumed from each electron donor, multiplied by 100 (Equation 6).

$$e - \ mol\ substrate = \ Consumed\ substrate(mol) \times Total\ available\ electrons\ in\ substrate \quad (4)$$

$$e - mol\ PHB = PHB\ as\ crotonic\ acid(mol)\ x\ electron\ required\ for\ crotonic\ acid\ synthesis \quad (5)$$

$$\% \, electron \, yield = 100 \times \frac{e{-}mol \, PHB}{e{-}mol \, substrate} \quad (6)$$
